# An Assessment of the Functional State of Endothelial Colony Forming Cells from Patients with Diabetes Mellitus and Chronic Limb Threatening Ischemia

**DOI:** 10.1101/2025.03.24.644359

**Authors:** Caomhán John Lyons, Michael Creane, Nadeem Soomro, Clara Sanz-Nogués, Lidia Shafik, Alicja Straszewicz, Tomás P Griffin, Alan Stitt, Timothy O’Brien

## Abstract

Chronic limb threatening ischemia (CLTI) is the most severe form of peripheral vascular disease which can lead to amputation with a high associated mortality rate. Endothelial colony forming cells (ECFCs) show potential as a cell therapy to revascularize the limbs of individuals with CLTI. However, autologous ECFCs from patient peripheral blood (PB) have been reported to have a dysfunctional phenotype. We investigated this disease phenotype in individuals with CLTI, with and without diabetes mellitus (DM), to determine ECFC suitability as an autologous cell therapy.

PB-ECFCs were isolated from age-matched controls, individuals with DM, and individuals with CLTI, with and without DM. The frequency of isolating ECFCs from this donor cohort was calculated. Furthermore, in vitro characterization assays were performed (growth kinetics, angiogenic properties, and reactive oxygen species (ROS) levels) and compared between donor groups.

We report a significantly increased frequency of ECFCs from individuals with CLTI, with and without DM. Furthermore, our results demonstrate no significant disease related effect on the *in vitro* functional properties of ECFCs between cohorts. However, there is a significantly higher *in vitro* angiogenic capacity in individuals with DM vs age-matched controls.

Our results demonstrate that ECFCs can be isolated in individuals with CLTI, with and without DM, and that ECFC functionality is similar between cohorts. Therefore, if the 70% isolation efficiency from CLTI cohorts is overcome, then autologous PB-ECFCs may be a suitable therapeutic for CLTI. Further analysis is needed to determine the critical quality attributes of ECFCs from this patient population.

**Significance Statement:** To the authors knowledge, this paper shows for the first time that endothelial colony forming cells can be isolated from individuals with chronic limb threatening ischemia, with and without diabetes. Additionally, we show a significantly higher frequency of endothelial colony forming cells isolated from chronic limb threatening ischemia patient cohorts. There is no significant difference in endothelial colony forming cells between age-matched controls and chronic limb threatening ischemia patient with and without diabetes mellitus in vitro, potentially suggesting an autologous approach may be a viable therapeutic option in the future.

## Introduction

Chronic limb threatening ischemia (CLTI) is the most severe form of peripheral artery disease and contributed to 3.9% of hospitalizations in the US in 2016-2019 [1]. CLTI is caused by the build-up of an atherosclerotic plaque in peripheral vessels causing ischemia. Individuals with CLTI experience pain at rest with the potential formation of ulcers. Individuals with CLTI have a poor prognosis, with an estimated 20% amputation rate within 5 years with a high associated mortality rate (48% within 5 years) [2,3]. Standard treatment for individuals suffering from CLTI is with either angioplasty or vascular bypass. However, 10-25% of individuals with CLTI are not suitable for this therapy and are considered ‘no-option’ and receive conservative management [4–6]. This ‘no-option’ cohort has a poor prognosis and often rapidly progresses to amputation. Diabetes mellitus (DM) is a major risk factor for CLTI and individuals with DM-CLTI rapidly progress to amputation. Consequentially, individuals with DM-CLTI have a higher amputation rate (34% within 5 yrs) and a higher mortality rate than individuals with CLTI alone [2]. A novel therapy for no-option individuals with and without DM would prevent amputation, reduce the mortality rate, and consequently reduce the healthcare and financial burden of CLTI [7].

Endothelial colony forming cells (ECFCs) are a potential therapeutic for this ‘no-option’ patient population. ECFCs are the true endothelial progenitor cell [8–10], and can directly form blood vessels *in vivo* and *in vitro.* ECFCs are characterized by having a cobblestone morphology, a robust proliferative potential, and the absence of the immune markers CD14 and CD45 [8–11]. ECFCs also release an array of pro-angiogenic factors such as angiogenin, endothelial growth factor, and endoglin, to support blood vessel formation [12,13]. Studies have shown the potential use of ECFCs in several *in vivo* models of ischemic conditions such as stroke [14,15], and ischemic retinopathy [16]. In hindlimb ischemia models (HLI) of CLTI, administered ECFCs can home to the site of injury and promote increased blood flow recovery, increased capillary density, and reduced tissue necrosis [17–19]. This indicates that ECFCs have potential as a therapeutic for ischemic conditions, however, there has been no clinical trial using ECFCs to date.

ECFCs can be isolated from both umbilical cord blood (UC) and peripheral blood (PB). From a therapeutic perspective, UC-ECFCs have a high proliferative, angiogenic and wound healing capacity [9,20], making them an ideal off-the-shelf therapeutic product for ischemic conditions. However, UC-ECFCs can trigger an immune response upon transplantation *via* MHC class I + MHC class II expression [21,22]. Therefore, translational studies have investigated the viability of an autologous PB-ECFCs therapy to treat vascular conditions. Studies have investigated the functionality of ECFCs from individuals with pulmonary arterial hypertension, von Willebrand’s disease, vascular thromboembolic disease, chronic obstructive pulmonary disease, and individuals with coronary artery disease [23–27]. The results from these studies showed that ECFCs from individuals with disease often present with an abnormal frequency and a dysfunctional phenotype, such as reduced tube formation, reduced migration, delayed colony formation, premature senescence, and a reduced ability to isolate ECFCs [23,26,28]. A disease related dysfunction has also been observed in ECFCs from individuals with DM such as reduced migration capacity, lower angiogenic potential, and slower proliferation [28–31]. This dysfunction has also been shown in *in vitro* models of hyperglycemia [32,33], and in ECFCs from gestational DM births [33–35]. To date, there have been no studies of ECFCs from individuals with CLTI with or without DM.

A disease related phenotype in autologous ECFCs would limit their therapeutic potential to treat individuals with CLTI. Therefore, we sought to assess the *in vitro* properties of ECFCs from individuals with CLTI, with and without DM, to identify, for the first time, whether ECFCs can be isolated from these patients and if any dysfunction exists in the patient populations. This would determine the suitability of autologous ECFCs as a therapy for CLTI and DM-CLTI.

## Materials and Methods

### Ethics and ECFC Isolation

Ethical approval was obtained from the Clinical Research Ethics Committee at University Hospital Galway (C.A. 1676). Age-matched controls (individuals with no DM or vascular disease), DM, and individuals with CLTI, with and without DM, were recruited (further details in Supplementary Materials). Patients were excluded based on inappropriate clinical characteristics (Figure 1). ECFCs were isolated from the PB using density centrifugation. ECFCs were cultured in EGM-2MV media (Lonza, CC-3202) with 16% fetal bovine serum (FBS), hereafter referred to as EGM. After 14 days in culture the number of colonies were counted and compared between donor groups. Experiments were conducted on ECFCs between passages 5-7.

**Figure 1.**
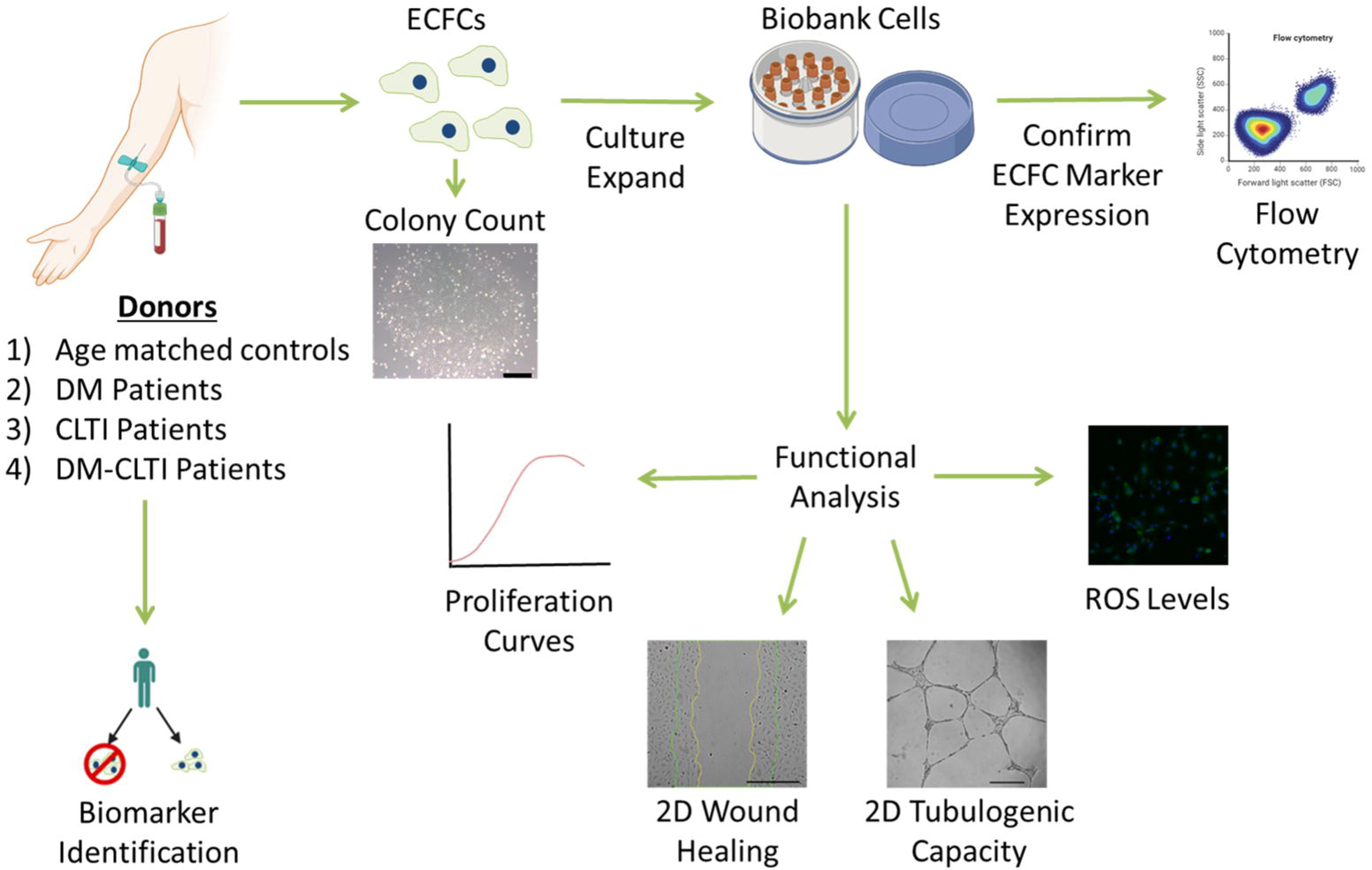
Flow chart of the patient exclusion criteria per patient group. Alt Text: Flow diagram of the patient exclusion criteria with patient numbers included.

### Immunophenotype Characterization

ECFC identity was confirmed using CD31^+^, VEGFR2^+^, CD34^low^, CD45^-^ (further information in Supplementary Materials).

### Proliferation Assay

To determine the proliferation capacity of ECFCs, cells were serially passaged at 6,000 cells/cm^2^ until replicative senescence and counted using Trypan blue (Sigma, T8154). The population doubling (PD) was calculated using Equation 1 and summed together to determine the maximum cumulative PD (CPD) level for each donor. To examine the proliferation rate, the PD time (PDT) was calculated between P3-5 as per Equation 2 and compared between donor cohorts [6,30].

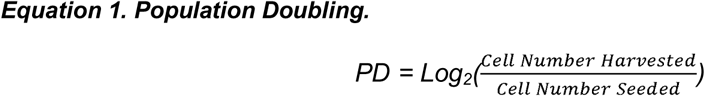

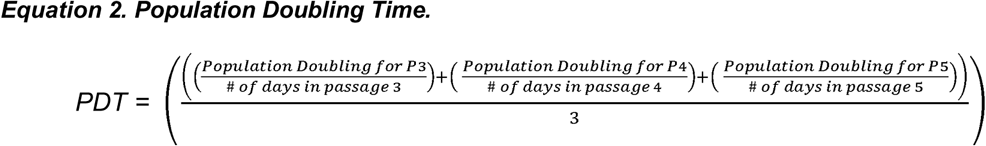

### 2D Wound Healing Assay

ECFCs were seeded into 48 well plates at 60,000 cells at P5 per well and left overnight to reach confluency. Each well was then scratched with a P200 pipette tip. The cells were gently washed with EGM and then fresh EGM was added to each well. Images were captured of the wound gap at 0 hr and at 10 hrs post scratching at 4X magnification using a Cytation 1 Cell Imaging Multi-Mode Reader (Agilent Biotek) with Gen5 software (v3.04). The % wound closure was calculated using Formula 3.

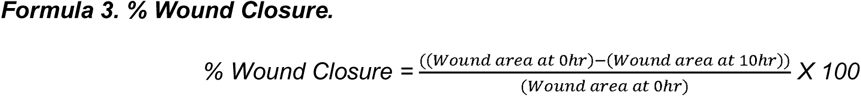

For media comparison experiments, the experiment was conducted as above except that cells were instead washed with basal media without growth factors or serum (EBM) prior to scratching to remove any residual EGM. After scratching the cells were cultured in either EGM or EGM-2 (Lonza, CC-3162), a media formulation which differs from our EGM by containing heparin sulphate and 2% FBS.

### In Vitro Matrigel Angiogenesis

A 48 well plate was coated with 110 μl of growth factor reduced Matrigel (Corning, 356231) for 1 hr at 37°C. 22,000 ECFCs at P5 were seeded dropwise per well in 500 µl of media. Cells were imaged at 4X with an Olympus CKX41 Upright Microscope after 18 hrs incubation at 37°C. The number of vessels, intersections (where two or more vessels connect), and complete loops were quantified using ImageJ (Version 2.14.0/1.54f).

For media comparison experiments 88,000 ECFCs from each donor were placed into two tubes containing EBM. Cells were centrifuged at 400 g for 5 min and the supernatant removed. Cells were then resuspended with either 2 ml of EGM or EGM-2 and seeded and assessed as above.

### Reactive Oxygen Species Analysis

To determine the level of reactive oxygen species (ROS), ECFCs were seeded at 6,000 cells/cm^2^ into two T25 flasks at P6. At ∼80% confluency the cells were washed with Hanks’ balanced salt solution (HBSS) (Thermofisher, 14025050). ECFCs were treated with 50 µM dichlorofluorescein-diacetate (DCFH-DA) (Merck, D6883) diluted in HBSS or with HBSS alone for 20 mins at 37°C in the dark. Cells were washed with PBS, passaged, centrifuged (all done at 400 g for 5 min), and then resuspended with HBSS. Cells from each flask (DCFH-DA treated and untreated) were equally split into two tubes, centrifuged, and resuspended with either HBSS alone or HBSS with 200 µM H_2_O_2_. Cells were incubated for 20 mins at room temperature in the dark. Cells were centrifuged, the supernatant removed, and then resuspended with 200 µl of HBSS. Cells were then treated with 2 nM Sytox far red viability dye (Thermofisher, S34859) in flow cytometry buffer for 7 mins to exclude dead auto fluorescent cells and run on a flow cytometer with 10,000 cell events recorded. The fluorescence level was normalized to cell size to account for differences in cell size affecting fluorescence levels and then measured as fold difference from the Sytox alone group (Equation 4).

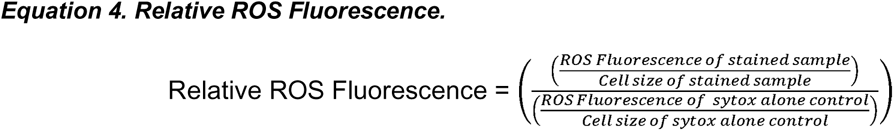

### Cell Metabolic Rate

To determine whether EGM masked any cell dysfunction, the metabolic rate of ECFCs cultured in EGM was compared to ECFCs cultured in EGM-2. ECFCs were seeded into a 96 well plate and incubated at 37°C overnight. Cells were then washed twice with EBM and then cultured with either EGM or EGM-2 for 48 hrs. The media was then removed from the wells and replaced with 110 µl of media containing PrestoBlue (Thermofisher, A13262) and incubated at 37°C, made as per manufacturers protocol. After 2 hrs, 50 µl of the PrestoBlue media from each well was plated into a 96 well plate in duplicate. The PrestoBlue fluorescence signal was measured at 572 nm for 0.1 s on a VICTOR^3^ Multilabel Plate Reader (Perkin Elmer) using the Wallac 1420 software. Cells were then stained with 8.1 µM Hoechst 3342 diluted in the respective media for 5 mins at room temperature before reading the fluorescence at 460 nm with a scan time of 0.1 s. The metabolic rate per cell was then calculated using Equation 5.

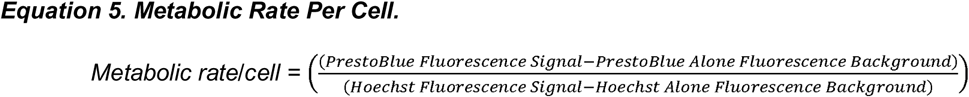

### ECFC Therapeutic Dose Estimation

An estimated therapeutic dose of ECFCs for therapeutic delivery to individuals with CLTI and DM-CLTI was determined using ECFC doses used in *in vivo* HLI mouse models from the literature [19,29,32,36–41]. The estimated therapeutic dose range was generated by taking the lowest and highest value from the literature, then scaled up by relative weight to reach a human dose. To calculate the ECFC yield we used the initial colony number and the cumulative population doubling until P7 (chosen as the upper passage threshold used by other studies [36,38,42]) (Equation 6). The cell number for release criteria were also included for an estimate of the therapeutic cell number needed for an ECFC therapy (Supplementary Table S1).

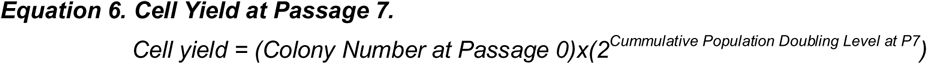

### Statistics

All data are represented as mean ± standard deviation (SD). Shapiro Wilk test was used to determine whether the sample data came from a normally distributed population. The differences between two groups with normal distribution were tested using an unpaired T-test. Differences between multiple normally distributed groups were tested statistically using one-way analysis of variance (ANOVA) with a Sidak post-hoc test. Analyses were performed using GraphPad Prism Version 9. For biomarker identification, patient clinical measurements were divided into two groups based on successful or failed ECFC isolation. A Student’s T-test or Mann-Whitney test was conducted as appropriate to identify any potential significance difference between these two groups. Binary logistic regression was performed to aid in the identification of potential predictors of ECFC isolation using the patient clinical data using Minitab version 19.2020.1.0. Significance was defined as P ≤ 0.05.

## Results

### Patients Were Matched for Age and Gender

Patient characteristics were noted upon recruitment to ensure that the patient groups were matched with regards to age and gender. From Table 1, the patients recruited in this study were matched based on age and gender. As expected, there was significantly higher HbA1c in individuals with DM and DM-CLTI (P<0.0001). With regards to the level of CLTI between patient cohorts, there was a significantly higher ABI in individuals with DM-CLTI compared to CLTI alone (P=0.04), however this is likely due to calcification of the vessel in individuals with DM-CLTI making it difficult to accurately measure the ABI [43].

**Table 1.** Patient characteristics. Mean ± SD. * = Significantly different from AMC. † = Significantly different from CLTI. $ = Significantly different from DM. Significant parameters are in bold. LDL was significant by one-way ANOVA but not significant by Sidak post-hoc test. ABI = Ankle-Brachial Index, ALT = Alanine Transanimase, aPTT = Activated Partial Thromboplastin Time, BMI = Body-Mass Index, eGFR = Estimated Glomerular Filtration Rate, HDL = High Density Lipoprotein, LDL = Low Density Lipoprotein, PT = Prothrombin Time.

From the patient blood biochemical parameters measured there was significantly higher cholesterol levels in age-matched controls vs individuals with DM (P=0.02), with similar statin use between age-matched controls and individuals with DM, indicating that the lower cholesterol levels may be due to the higher use of biguanides in the DM group which has been shown in the literature to reduce cholesterol levels [44]. With regards random glucose levels, only individuals with DM-CLTI had significantly higher blood glucose levels compared to age-matched controls (P=0.04). Additionally, as expected, use of NSAIDs, biguanides and statins are higher in their respective disease cohorts (P=0.02, 0.003, 0.01 respectively).

### Significantly More ECFC Colonies in Individuals with CLTI and DM-CLTI Compared to Age-Matched Controls

The literature has reported that the frequency and the ability to successfully isolate ECFCs from a variety of disease cohorts can be altered with reduced levels usually being reported [35,45]. We compared the isolation efficiency in our patient cohorts relative to the Age-Matched Controls to determine whether any patient cohort had a reduced ability to isolate PB-ECFCs (Figure 2). The criteria to determine a successful isolation was determined based on the ability to culture expand the cells and confirmation of the ECFC phenotype (CD31^+^, VEGFR2^+^, CD34^low^, CD45^-^)(Supplementary Figure S1). In this study we noted no significant difference in the isolation rate between cohorts (Figure 2A), suggesting that DM and CLTI are not significantly impacting ECFC isolation rate. We report that ECFCs can be isolated from both CLTI cohorts, with an isolation rate of 69.6%.

**Figure 2.**
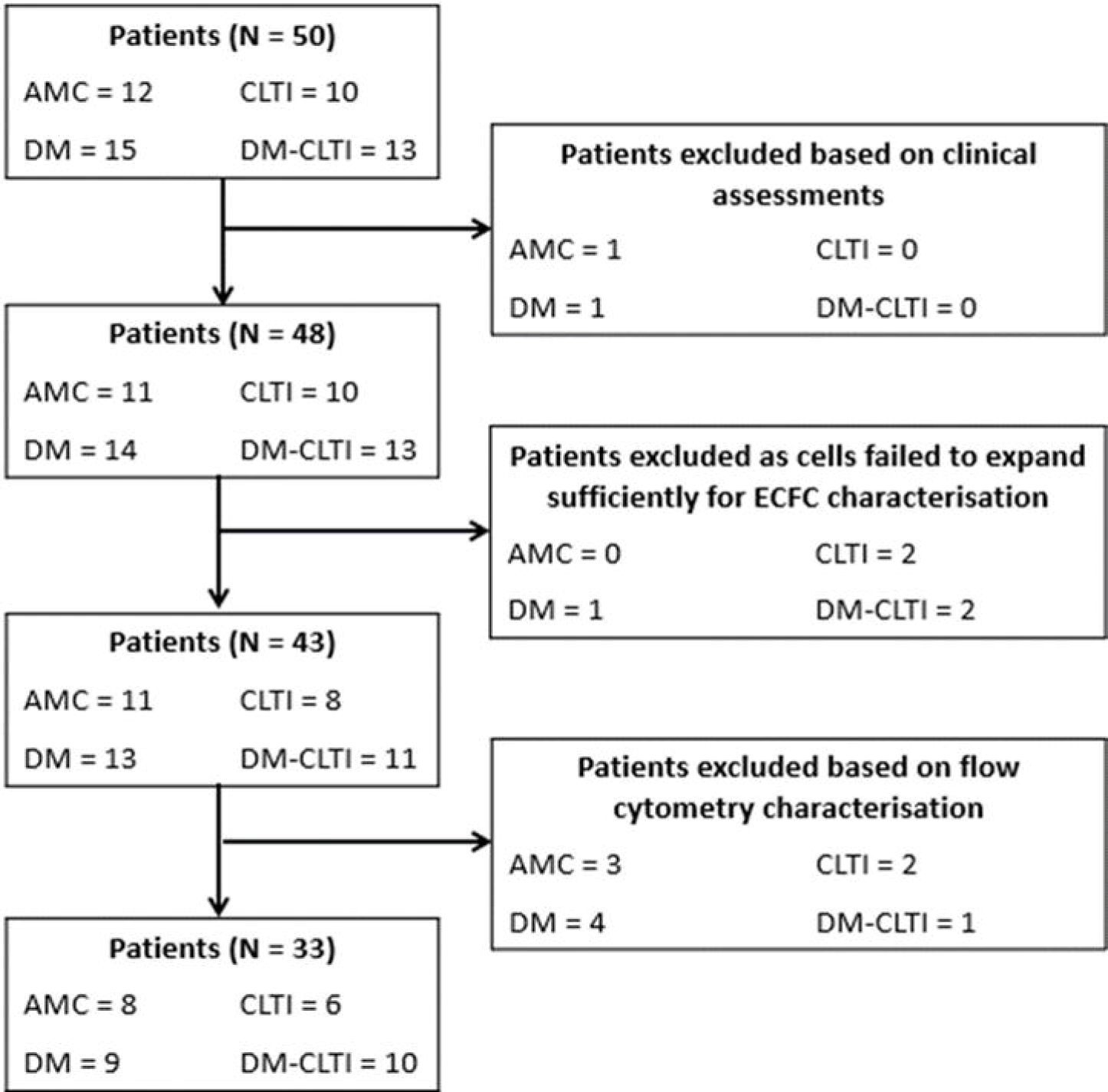
Significantly higher frequency of ECFCs from the PB of individuals with CLTI and DM-CLTI vs AMCs. A) ECFC isolation efficiency from each patient cohort is noted above each pair of bars. There was no significant difference in the isolation efficiency between the donor groups (P = 0.71). B) Colony number after two weeks in culture showing a significantly higher number of colonies in both the CLTI and DM-CLTI patient cohorts compared to age-matched controls (AMCs) (P = 0.023 and 0.047 respectively). Hollow points = Female Donors, Solid points = Male Donors. C-J) ECFC colony counts at two weeks divided by drug class use. There was a significantly higher ECFC colony number in individuals with β-blockers than those without β-blockers (P = 0.008), but no significant difference between individuals with or without biguanides, sodium-glucose co-transporter-2 (SGLT2) inhibitors, angiotensin-converting enzyme (ACE) inhibitors, Factor Xa inhibitors, statins, non-steroidal anti-inflammatory drugs (NSAIDs) and proton pump inhibitors (PPIs) (P = 0.45, = 0.16, = 0.93, = 0.05, = 0.09, = 0.40, = 0.49, respectively). K) Representative images of ECFC colonies (*) after two weeks from each patient cohort. Scale bar = 500 µm. Mean ± SD. * p = < 0.05, ** = < 0.01. **Alt Text:** A series of graphs demonstrating ECFC isolation efficiency and the number of colonies isolated from patients from each cohort, also broken down by drug use. Representative images of colonies from each donor cohort are included.

To determine whether the frequency of ECFCs was altered in the disease patient cohorts, the number of ECFC colonies formed after two weeks culture from each patient cohort was quantified (Figure 2B). We demonstrate that there was a significantly higher number of ECFC colonies in DM-CLTI donors (P=0.047) and CLTI donors (P=0.023) compared to Age-Matched Controls, with no gross morphological differences observed in the colonies between patient cohorts (Figure 2K).

As part of best medical therapy all recruited patients were all kept on their respective medication. As a result, we sought to investigate if the medication impacted the frequency of ECFCs isolated. The drugs that each patient was taking at blood draw were noted and then grouped based on drug class. The patients were then divided into those who received that drug class vs those who did not. The results show that only patients who received β-blockers had significantly higher number of ECFC colonies than those who did not receive β-blockers (P=0.008) (Figure 2C-J).

### Patient-Derived ECFCs Have a Similar Expansion Capability as ECFCs From Age-Matched Controls

Disease environments can dramatically impact the expansion capability of cells which can be detrimental for an autologous therapeutic approach. To determine whether DM or CLTI reduced the growth kinetics of ECFCs from the patient cohorts, cells were cultured until replicative senescence. We observed no significant difference in the growth kinetics of ECFCs between the donor groups (PDT P=0.26, and maximum CPD P=0.18) (Figure 3A-C). This data suggests that ECFCs from patient groups have similar growth kinetics as those derived from Age-Matched Controls.

**Figure 3.**
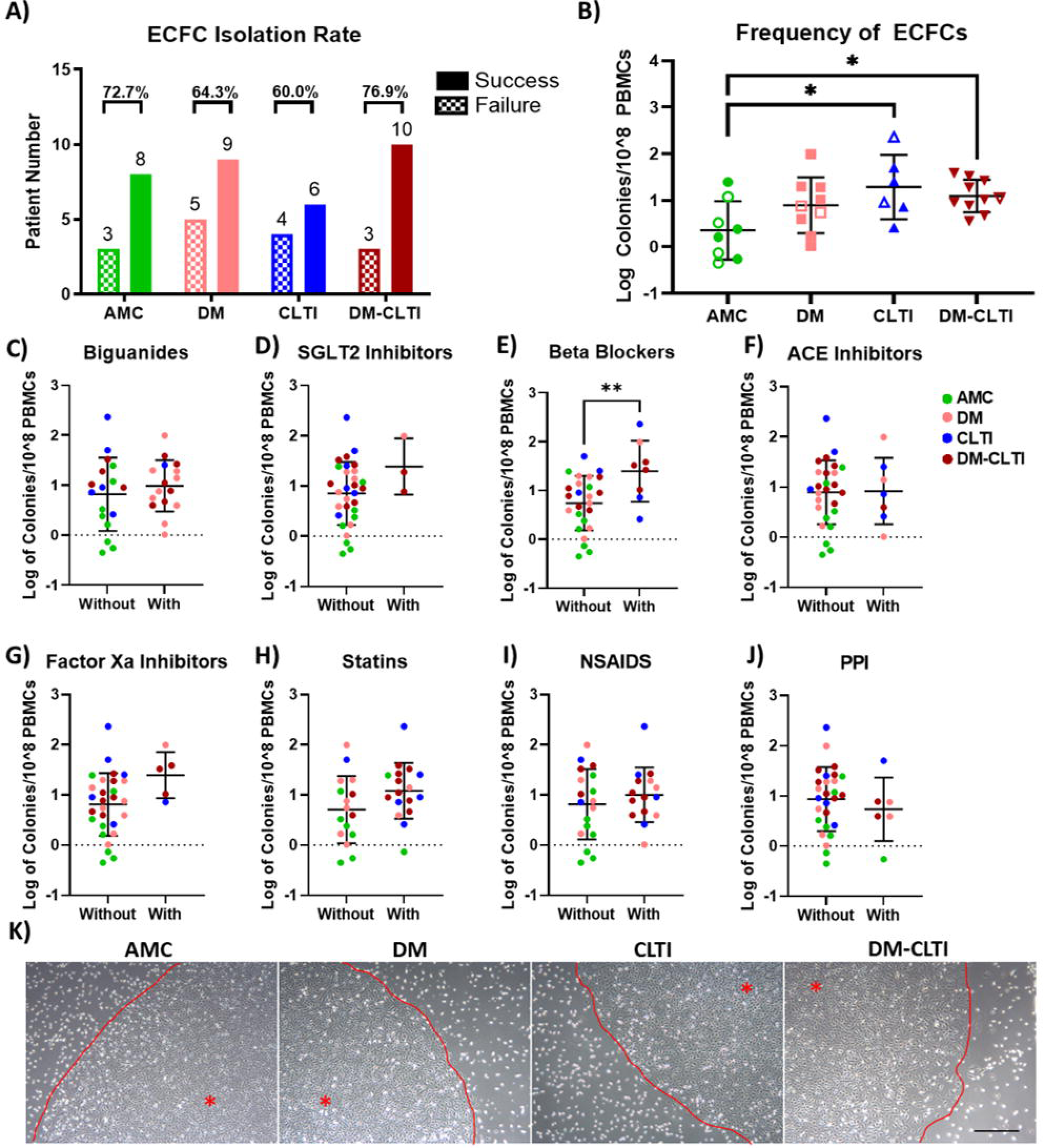
No significant difference in the growth kinetics of ECFCs between patient cohorts with a high proportion of donors yielding sufficient ECFCs for an autologous therapy at the minimum therapeutic cell number. A) Cumulative population doubling of ECFCs from all donor groups. B) Maximum cumulative population doubling capacity of ECFCs with no significant difference observed between the four donor groups (P = 0.11). C) The average population doubling time of ECFCs between passage 3 – 5 (P = 0.09). Hollow points = Female Donors, Solid points = Male Donors. D) The PB-ECFC yield from patients was calculated at P7 and an estimated therapeutic dose was calculated using ECFC doses from HLI studies in the literature. The estimated therapeutic dose was then scaled by relative weights between mouse and patient weights. The cell number for release was then added to get the therapeutic cell number. & = no weight was recorded for this patient, so the estimated therapeutic dose was scaled based on the average weight within the CLTI group. Mean ± SD. * p = < 0.05. **Alt Text:** Graphs of the proliferative properties of ECFCs between the patient cohorts.

With the main aim of this work being to evaluate the potential of an autologous ECFC therapeutic, one of the key questions to address is whether PB-ECFCs from individuals with CLTI, with and without DM, can be expanded sufficiently to yield a therapeutic dose. We plotted the ECFC yield (at P7) from each patient and used an estimated therapeutic dose based on the minimum and maximum therapeutic dose in animal models of HLI in the literature. The estimated therapeutic dose was then scaled up to the level of each patient using their weights. Only 2/7 Age-Matched Control donors reached the minimal therapeutic cell number; in contrast, a higher proportion of individuals from disease cohorts reached the minimum therapeutic cell number (4/6 DM, 6/6 CLI, 7/8 DM-CLI) (Figure 3D). Based on this result an autologous PB-ECFC therapeutic for individuals with CLI, with and without DM, may be suitable; however, it should be noted it took an average of 38.03 7.12 days for ECFCs in patient samples to reach P7.

### No Significant Difference in the In Vitro Angiogenic and Wound Healing Capacity of ECFCs from CLTI Patient Cohorts

As the angiogenic capacity and the ability for cells to migrate towards damage are two of the principal functions of ECFCs we assayed the *in vitro* tubulogenic capacity of the ECFCs, using a 2D Matrigel tube formation assay, and the wound healing capacity, using a scratch assay. After incubating 18 hrs there were no significant difference in the tubulogenic capacity of ECFCs from individuals with CLTI and with DM-CLTI, however there was a significantly higher number of tubules (P=0.02) and intersections (P=0.03) between ECFCs from individuals with DM vs age-matched controls (Figure 4A-D). Despite the altered angiogenic capacity there was no significant difference in the wound closure rate after 10 hrs compared with control donors (P=0.51) (Figure 4E+F). The above data indicates that there is increased tubulogenic capacity of ECFCs from individuals with DM compared to Age-Matched Control ECFCs and this effect is diminished to the level of non-significance in individuals with DM-CLTI, with no apparent effect of DM or CLTI on the ability for ECFCs to close wounds *in vitro*.

**Figure 4.**
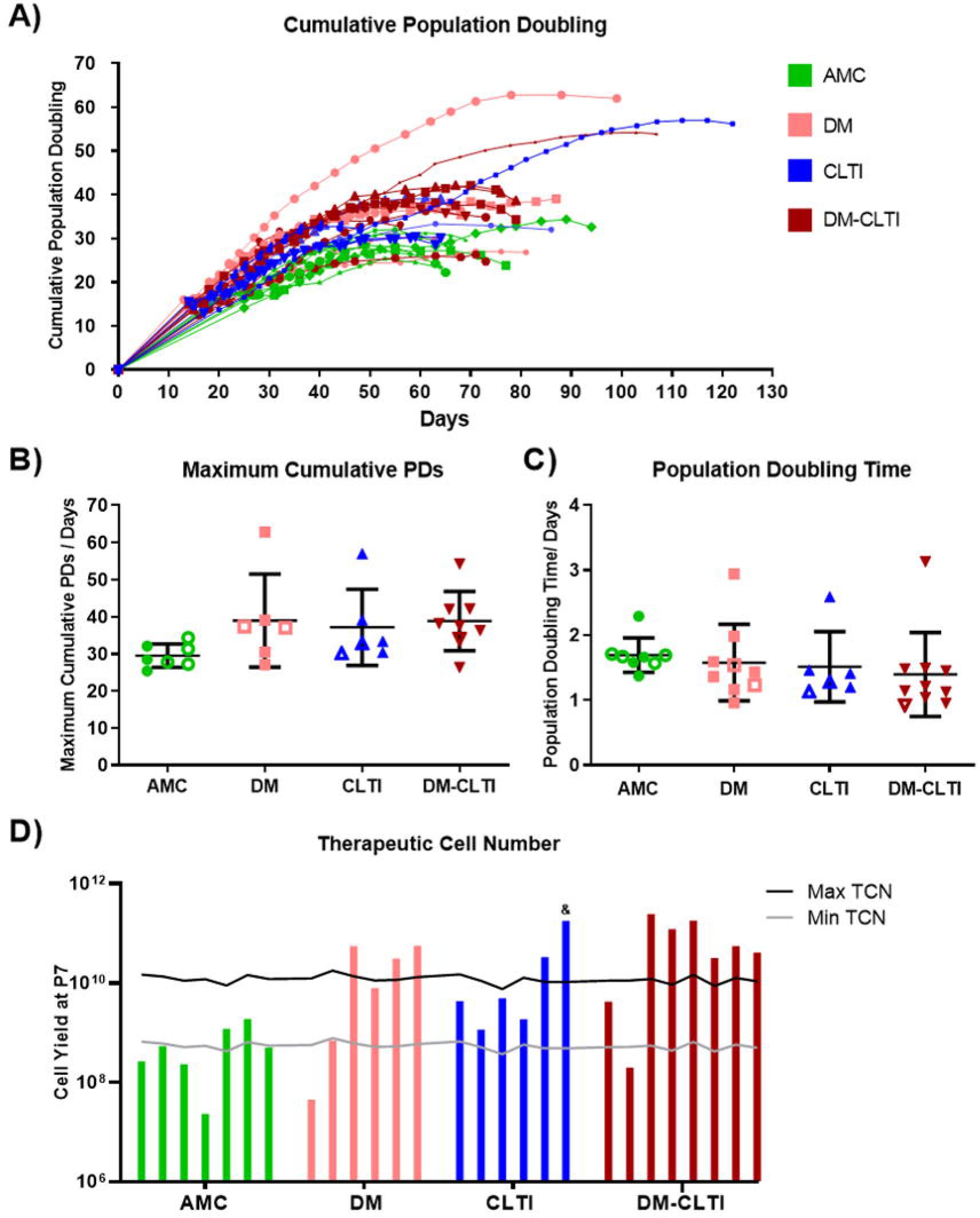
Significantly higher tubulogenic capacity in ECFCs from individuals with Diabetes Mellitus vs Age Matched Controls however no change in wound closure rates or ROS levels. A) Representative images of tube formation in the different donor groups after 18 hrs incubation on growth factor reduced Matrigel. Number of tubes (B), closed loops (C), and intersections (D), defined as the connection of two or more tubes, were quantified. There was a significantly higher number of tubes and intersections in individuals with DM compared to AMCs (P = 0.02 and 0.03 respectively). There was no significant difference in the number of closed loops between groups (P = 0.17). E) Representative images of the wound closure per group. F) Wound closure percentage between the donor groups indicating that there was no significant difference in the wound closure rate of ECFCs from different donor groups (P = 0.51). G) Gating strategy for ROS staining. H) Resting intracellular ROS levels demonstrating no significant difference at baseline levels between groups (P = 0.63). I) Intracellular ROS levels after H_2_O_2_ stimulation with no difference between groups in the response to a ROS inducing treatment (P = 0.74). Mean ± SD. * p = < 0.05. Scale bar = 500 µm. **Alt Text:** Series of graphs plotting the angiogenic and wound healing capacity of ECFCs with representative images. Graphs of the ROS levels in ECFCs are also included with the flow gating strategy used.

### The ROS Levels were not Significantly Different Between Donor Groups

ROS is an important chemical mediator for angiogenic function; however, elevated levels can result in cell damage and apoptosis. In DM and in CLTI the levels of ROS are much higher than in the normal healthy environment [46–50]. Therefore, we sought to identify whether there were higher endogenous ROS levels in ECFCs from the patient cohorts vs Age-Matched Controls. ECFCs were incubated with DCFH-DA and the level of fluorescence was measured using flow cytometry. The gating strategy used is indicated in Figure 4G. We also measured the levels of intracellular ROS in ECFCs in response to H_2_O_2_ treatment to determine whether ECFCs from patient cohorts would have a different response to ROS insult compared to ECFCs from Age-Matched Controls. We found no significant difference in the endogenous and stimulated levels of intracellular ROS between the donors (P=0.63, =0.74 respectively)(Figure 4H+I).

### EGM Did Not Compensate for Disease Related Dysfunction

Despite our initial hypothesis that ECFCs from individuals with DM and CLTI would display a dysfunctional phenotype *in vitro*, our results show minimal disease phenotype in ECFCs from the different patient cohorts. It should be noted, however, that these assays are performed after a period in culture which may alter the cell phenotype compared to the native uncultured ECFC. This study used a media with a high FBS concentration to improve ECFC isolation, therefore we sought to ascertain whether this FBS high media compensated for any disease related dysfunction. To assess this, we compared the effect of our media (EGM) on ECFCs against EGM-2, a more commonly used ECFC media formulation with lower serum levels. We compared the metabolic rate, 2D wound healing capacity, and 2D tubulogenic capacity of ECFCs cultured using both EGM and EGM-2 to identify whether a disease phenotype would be present in the EGM-2 and not in the EGM cultured ECFCs.

ECFCs were initially cultured in EGM and then for each assay the cells were washed with EBM before being cultured in either EGM-2 or EGM. There was no significant difference in the metabolic rate (P=0.12), wound healing capacity (P=0.79), or tubulogenic capacity (number of tubes P=0.03 (not significant by Sidak post-hoc test), number of intersections P=0.12, number of loops P=0.42) between the donors within either media group (Figure 5). This suggests that the EGM media used in this study did not compensate for any disease related dysfunction associated with the angiogenic capacity of ECFCs from the diseased patient groups.

**Figure 5.**
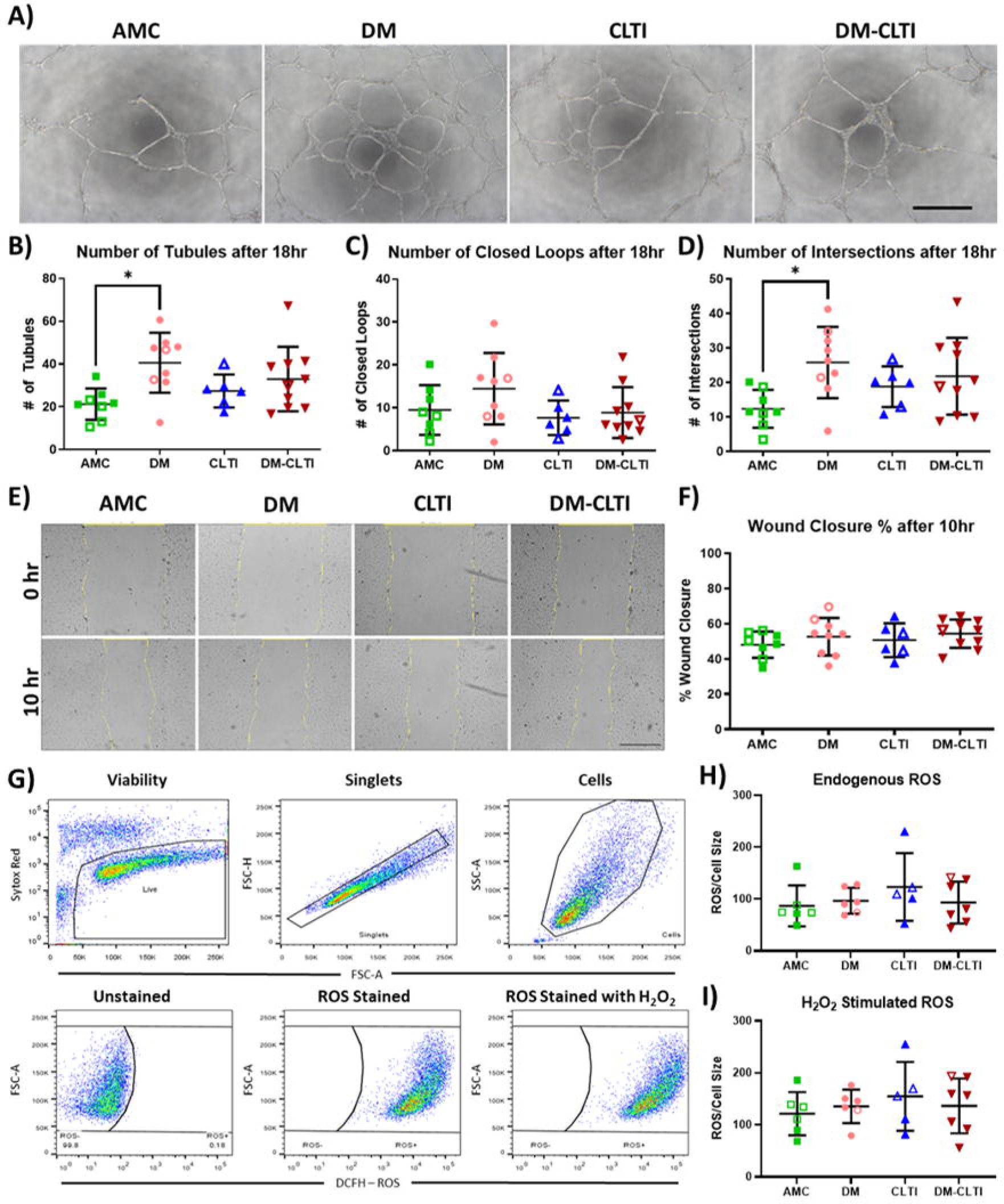
A higher serum concentration does not mask disease related dysfunction in patient derived ECFCs. A) ECFCs were cultured with either EGM (the media used in this study) or EGM-2. The metabolic rate was measured by PrestoBlue and normalised to the number of cells using Hoechst fluorescence. B) The % wound closure in both EGM and EGM-2 was measured after 10 hours. C-F) Quantification of the tube network formed in the in vitro Matrigel assay after 18 hours. There was no dysfunction observed in the above parameters between the donor groups in either media group. Solid points = Male, Hollow points = Female. Mean ± SD. *p = < 0.05. Scale bar = 500 µm. **Alt Text:** Graphs and representative images of the angiogenic capacity of ECFCs, with graphs on the metabolic activity and wound closure ability of ECFCs, demonstrating no functional difference between donor cohorts with two different media formulations.

### ECFC Colony Number and Systolic Blood Pressure Correlate with Successful ECFC Isolation Rates

To identify potential predictors of successful ECFC isolation, we examined relevant biochemical and physiological parameters for each outcome. We compared the difference in mean values of clinical parameters from individuals with successful ECFC isolation vs failed ECFC isolation, and results showed statistically significant differences in ECFC frequency, high density lipoprotein (HDL), creatinine, weight, mean corpuscular hemoglobin, mean corpuscular volume, eosinophil number, and systolic pressure (p<0.05) (Supplementary Table S2). We then investigated whether relevant clinical parameters, e.g., parameters relevant to DM and CLTI, had a significant effect on the odds of successful ECFC isolation using a univariate binary logistic regression (Table 2, Step 1). This model identified creatinine, ECFC frequency, HDL, and systolic pressure (p<0.05) as variables that had a significant relationship with the odds of successful ECFC isolation. Taking the significant parameters, we then fitted a multivariate logistic model (Table 2, Step 2) which identified ECFC colony number as the only useful predictor of ECFC isolation (P=0.044). To further refine the model, we included ECFC colony number and systolic blood pressure (Table 2, step 3) which explained ∼68% of the outcome variability, and identified both variables as useful predictors of ECFC isolation (p<0.05). Our refined model shows that the odds of having successful ECFC isolation significantly increased with increasing ECFC frequency and systolic blood pressure.

**Table 2.** Binary logistic regression model of ECFC isolation vs patients’ clinical parameters. Vales in bold are significant (P < 0.05). Adj = Adjusted, BMI = Body Mass Index, CI = Confidence Interval, HDL = High Density Lipoprotein, LDL = Low Density Lipoprotein.

## Discussion

The work presented here focused on identifying the dysfunctional phenotype between PB-ECFCs from individuals with DM and with CLTI, with and without DM. Our data demonstrates a significantly higher frequency of ECFCs from individuals with CLTI and DM-CLTI compared to Age-Matched Controls, with no difference in colony number between Age-Matched Controls and individuals with DM (Figure 2). While the frequency of PB-ECFCs from individuals with DM has not been compared in the literature, there are studies which have shown that UC-ECFCs from gestational diabetic pregnancies and in *in vitro* hyperglycemia models result in a lower number of colonies [33,35]. Furthermore, studies show that ECFCs from severe atherosclerotic patients have a lower number of ECFCs, whereas individuals with venous thromboembolic disease resulted in a significantly increased ECFC frequency [25,51]. When the effect of hypoxia (1% O_2_) on ECFCs was examined *in vitro* there was significantly lower ECFC colony appearance [52], therefore suggesting that hypoxia increases ECFC frequency via increased mobilization rather than through increased proliferation. This mobilization hypothesis is supported by the literature which demonstrates that ECFCs can migrate to hypoxic sites when given therapeutically [14,15]. Additionally, we report a significantly higher number of colonies with the use of β-blockers which also may be a contributing factor to the increased ECFC frequency. Besnier *et al* found no significant difference in ECFC frequency in patients with β-blockers, however they isolated ECFCs on gelatin coated flasks vs the collagen coated flasks used here [27]. Further research is needed to validate the effect of β-blockers on ECFC frequency.

This study reports an average isolation efficiency of ∼69%. Reports indicate an isolation rate of 21-90% in ECFCs from healthy control donors which is in keeping with the 73% isolation rate from our Age-Matched Control patient cohort [27,53–56]. Looking at the DM cohort, Jarajapu *et al* noted only an isolation rate of 15% from individuals with DM compared to the 64% isolation rate from individuals with DM reported here [56]. With regards to the CLTI cohorts, to date no other study has isolated ECFCs from individuals with CLTI; however, isolation efficiency of 33% and 13% respectively, versus the ∼70% isolation efficiency in studies examining systemic atherosclerosis and acute coronary syndrome patients show an CLTI cohorts in our study [51,57]. Interestingly, when ECFCs were successfully isolated from patients in the CLTI cohorts, they reliably reached a cell number sufficiently high for a proposed therapy (13/14 donors) vs only 2/7 donors in Age-Matched Controls. The higher isolation rate from disease patient cohorts compared to that reported in the literature may be due to the higher serum concentration to improve ECFC isolation rates, as recommended by Smadja *et al*. [53]. Overall, papers do not report the isolation efficiency of ECFCs from donors which may lead to the impression that ECFCs can be reproducibly isolated from all patient donors. The isolation efficiency of ECFCs is a clinically relevant parameter that demonstrates a hindrance on the path to a potential autologous therapy that should be reported [53].

Our data demonstrates that an increased ECFC frequency and an increased systolic blood pressure were significant predictors of a higher chance of successful ECFC isolation. While it seems logical that a higher ECFC frequency would lead to an improved capability to successfully culture ECFCs, the effect of systolic blood pressure on the success of ECFC isolation is not so clear. As high shear stress decreases endothelial cell proliferation [58], instead, we hypothesize this could be caused by increased mobilization of ECFCs as high shear stress increases endothelial cell migration [59,60]. Our result is supported by the literature which has previously shown that hypertension leads to improved isolation rates of ECFCs [27]. Besnier *et al* also showed that an increase in BMI >30 kg/m2 resulted in lower ECFC isolation efficiency which was not observed in our study (p=0.07); however, an increased number of patients could result in sufficient power to detect a significant difference. Future studies can validate these biomarkers for work using PB-ECFCs.

Our study demonstrates minimal in vitro functional differences between the patient groups and the Age-Matched Controls with regards to their proliferation, wound healing capacity, tubulogenic capacity, and ROS levels (Figures 3,4). Instead of a disease related dysfunction, ECFCs from individuals with DM had a higher tube formation and intersection number compared to ECFCs from Age-Matched Controls, suggesting an increased angiogenic capacity in ECFCs from individuals with DM (Figure 4). This result contrasts with what was previously reported in the literature which showed ECFC dysfunction from both individuals with DM and in *in vitro* hyperglycemia models which showed delayed colony formation, decreased angiogenic capacity, and decreased migration capacity [28,29,33,46,61]. Interestingly, in individuals with coronary artery disease there was significantly increased angiogenic function, however *in vitro* models of hypoxia reports have shown reduced colony formation, reduced angiogenic function, and increased apoptosis [8,27,52,62,63]. This disparity indicates that further research is needed to examine the disease related effects from CLTI on ECFCs. We report superior growth kinetics in our Age-Matched Control group than that reported in the literature (CPDs = 10-20 vs our 29.53, and PDT = ∼3.5-4 vs our 1.8 days) which may have been due to the higher serum used in this study [9,64]. Previous studies have not examined the ROS levels of ECFCs from individuals with DM or CLTI, however significantly higher ROS levels were reported in ECFCs from individuals with venous thromboembolic disease [25].

The difference in the ECFC *in vitro* characteristics between these studies and the data presented here may be due to several reasons. 1) The patients recruited in this study were all kept on medications which may have minimized any disease related dysfunction within the cells. 2) Differences in patient severity, e.g. the DM donors used in this study were relatively well controlled from a glycemic perspective compared to that which is reported in other studies and our Age-Matched Control cohort were less healthy than those reported in other studies (60-135 mmol/mol in the Langford-Smith *et al* study [28] vs 48-85 mmol/mol in our DM cohorts, and higher BMI in our age-matched control cohort vs the non-diabetic cohort in the study by Leicht *et al* [30]). While no study has been conducted on ECFCs from individuals with CLTI specifically, a study by Simoncini *et al* examined individuals with atherosclerotic cardiovascular disease [51]. The individuals recruited in their study were of similar age to those in this study, but they reported more male participants in the atherosclerotic cohort vs their non-disease control cohort. This is important as females with CLTI have a higher treatment related complication rate and poorer outcomes, which may suggest gender differences in CLTI pathophysiology [65–67].

3) The cells may be dysfunctional *in vivo*; however, when introduced into an *in vitro* healthy environment the normal functionality of the cells may be restored due to the absence of the ischemic and hyperglycemic environment. Previous studies have shown that a disease related environment can cause cellular dysfunction in healthy cells [8,33,46,52,61–63]; however, the removal of these stimuli may allow the cell to recover to a level of relative normal cell functionality. Future work in this study would necessitate examining the ECFCs in disease related environments such as an inflammatory, hyperglycemic, and/or hypoxic conditions. 4) Our study used a high serum concentration (∼16%) which is contrast to the lower (2-5%) serum which is used in other studies [9,17,27,57]. This high serum may have provided optimal conditions which adjusted for the disease related dysfunction. While it was a short-term study, we observed no disease related phenotype using a less FBS rich media formulation (EGM-2) vs our EGM, suggesting that media did not correct any disease related dysfunction. A limitation of this work is that all ECFCs were isolated with high serum to facilitate superior ECFC isolation. Isolating with high serum concentrations may correct the ECFC ‘dysfunction’ early in culture. 5) Furthermore, many studies examined ECFCs from patients at earlier passages (∼P3) [28,29,51] compared to that analyzed here (P5-7). Potentially the time in culture dampened the disease related phenotype to the point of no significant difference. To investigate this, future work could compare the *in vitro* functionality at P3 to determine whether the disease related dysfunction exists at an earlier passage. The above points indicate several possible reasons why we reported no significant dysfunction between our patient cohorts in contrast to the literature.

## Conclusion

In this study we have shown that ECFCs can be isolated from all ECFC patient cohorts stated previously with equal isolation efficiency and that ECFCs from individuals with CLTI, with and without DM, resulted in a significantly higher frequency in PB. Once expanded, ECFCs from the patient CLTI cohorts have similar *in vitro* function. The results from this work indicate that autologous PB-ECFCs from individuals with CLTI may be a potential therapeutic in the future. However, further research will be required to obtain an improved understanding of the critical attributes of ECFCs isolated from this patient population.

## Supporting information

Supplementary Methods + Results

## Abbreviations

ACE: Angiotensin-Converting Enzyme
ALT: Alanine Transanimase
AMC: Age Matched Controls
aPTT: Activated Partial Thromboplastin Time
BMI: Body Mass Index
CPD: Cumulative Population Doubling
CLTI: Chronic Limb Threatening Ischemia
DCFH-DA: Dichlorofluorescein-Diacetate
DM: Diabetes Mellitus
ECFCs: Endothelial Colony Forming Cells
eGFR: Estimated Glomerular Filtration Rate
FBS: Fetal Bovine Serum
FCB: Flow Cytometry Buffer
FMO: Fluorescence Minus One
FSC: Forward Scatter
HBSS: Hanks’ Balanced Salt Solution
HDL: High Density Lipoprotein
LDL: Low Density Lipoprotein
MFI: Median Fluorescence Intensity
NSAIDs: Non-Steroidal Anti-Inflammatory Drugs
PB-ECFCs: Peripheral Blood ECFCs
PBMCs: Peripheral Blood Mononuclear Cells
PBS: Phosphate Buffered Saline
PD: Population Doubling
PDT: Population Doubling Time
PPI: Proton Pump Inhibitors
PT: Prothrombin Time
ROS: Reactive Oxygen Species
SD: Standard Deviation
SGLT2: Sodium-Glucose Transport Protein 2
SSC: Side Scatter
UC-ECFCs: Umbilical Cord-ECFCs
VEGFR2: Vascular Endothelial Growth Factor Receptor 2

## Acknowledgements

Thanks to Dr. Linda Howard and Dr. Cynthia Coleman for their support and direction on the project. Thanks to Veronica McInerney and the clinical research facility for aiding with patient sample collection and ethics writing.

## Competing Interests

TOB is a founder, director, and equity holder in Orbsen Therapeutics Ltd.

## Availability of Data and Materials

Data can be made available upon reasonable request to the corresponding author.

## Funding

This work was supported by the Science Foundation Ireland (SFI) Investigator Program (15/IA/3136) and the SFI Frontiers for the Future Programme (20/FFP-A/8794).

---

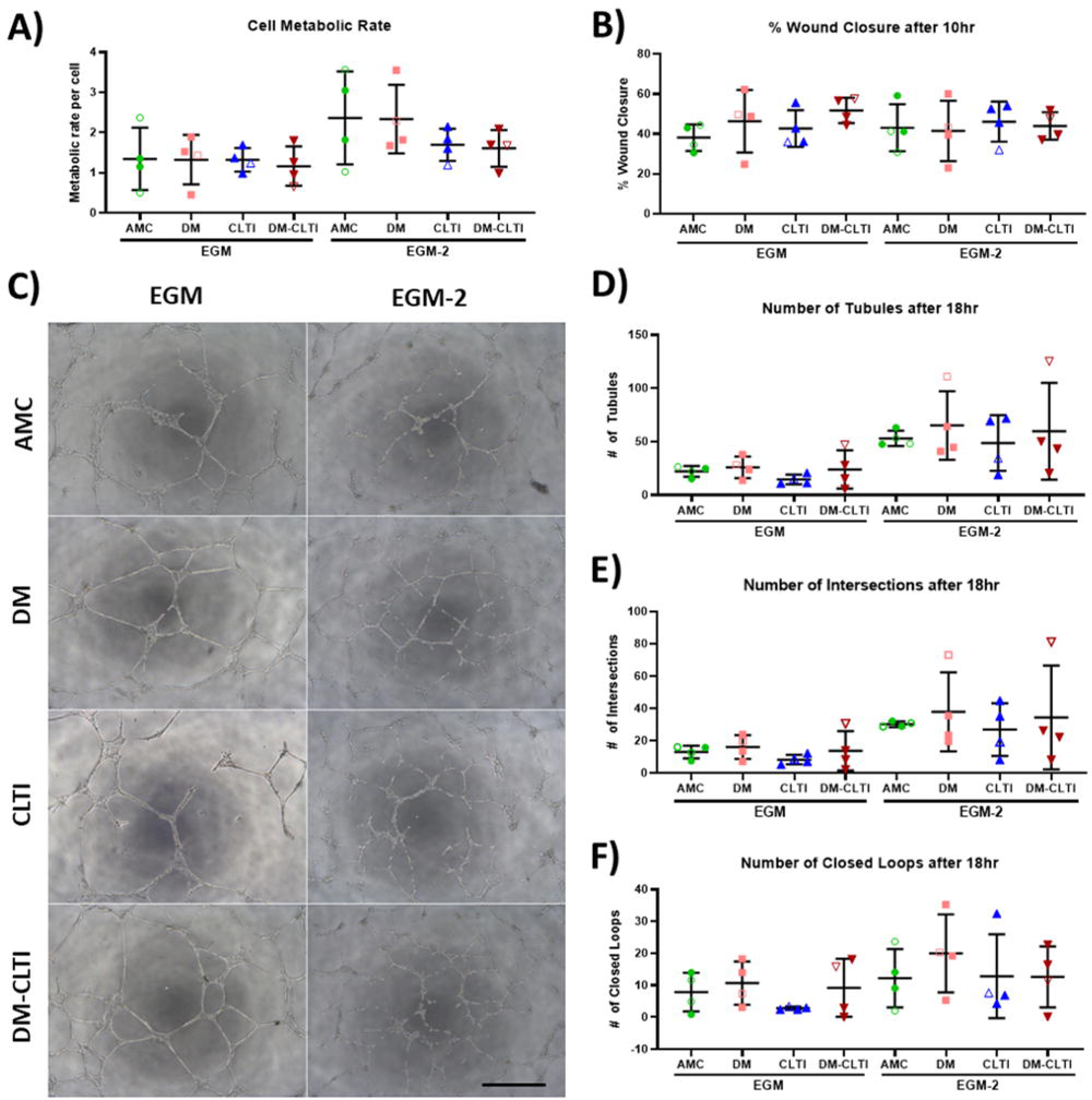

## Notes

**Acknowledgement** This publication has emanated from the research supported by Science Foundation Ireland (SFI) Investigator Program (15/IA/3136) and the SFI Frontiers for the Future Programme (20/FFP-A/8794). The authors acknowledge the facilities and the scientific and technical assistance of the University of Galway Flow Cytometry Core Facility.

