## Supplementary Methods + Results for "An Assessment of the Functional State of Endothelial Colony Forming Cells from Patients with Diabetes Mellitus and Chronic Limb Threatening Ischemia"

### Materials and Methods

#### Patients and Ethics

Adult peripheral blood (PB) was acquired from patients of University Hospital Galway with approval from the institutional review board (C.A. 1676) and in accordance with the Declaration of Helsinki. Age matched controls (AMCs) (defined as aged matched individuals with no diabetes or cardiovascular disease), patients with type 2 diabetes mellitus (DM), and patients with chronic limb threatening ischemia (CLTI), with and without DM were recruited. Written consent was acquired from each patient prior to a medical examination and a blood draw. A patient was defined as suffering from CLTI when they presented with a Rutherford score ≥ 4. Patients with DM were defined as suffering from DM based on having a HbA1c value >48 mmol/mol in their medical history. Patients were excluded if they presented with any hematological disorder, active infection, active malignancy or were unable to consent. 50 patients were recruited and all samples were processed within 2 hrs of blood draw. Blood was also sent to the hematology lab in University Hospital Galway for biochemistry analysis. While peripheral blood mononuclear cells (PBMCs) were isolated from each donor, several donors were excluded from the study for reasons such as: inappropriate clinical characteristics, failure to expand the cells and due to inappropriate flow cytometry cell phenotype (Supplementary Figure S1). These restriction criteria resulted in a drop in patient number from 50 to 33.

#### ECFC Isolation and Culture

To isolate endothelial colony forming cells (ECFCs) $\sim$80 ml of patient’s blood was harvested in lithium heparin tubes and diluted 1:1 with PBS. This was then carefully overlaid on Ficoll-Hypaque (GE-Healthcare, GE17-1440-02) at a ratio of 3:2 (3 parts blood/PBS mix:2 parts Ficoll-Hypaque). The tubes were centrifuged at 400 g for 35 mins (brake = 0) and then the PBMCs were carefully harvested from the buffy coat layer using a Pasteur pipette. The isolated PBMCs were diluted 1:1 with PBS and centrifuged at 300 g for 20 mins (brake = 9). The supernatant was then removed, and the cells were re-suspended with 6 ml of EGM-2MV (Lonza, CC-3202) (made without the accompanying FBS but instead mixed 5:1 with FBS (SV30160.03, GE Healthcare Hyclone)), this media is hereafter referred to as EGM. A T75 flask was coated with 50 μg/ml of rat tail collagen type I (Corning, 354236) (diluted in 0.02 M acetic acid and filter sterilized) for 1 hr at room temperature. The collagen solution was removed and washed with EGM to neutralize the acidic environment and to remove excess collagen-acetic acid solution. A cell count was carried out using Turk’s solution (Sigma, 1.09277). The cells were then plated into the collagen coated flask and placed in a standard cell culture incubator at 37^o^C with 5% CO_2_. After 24 hrs the media was changed and was subsequently changed every other day until colonies were sufficiently large to passage with trypsin. A colony count was performed on day 14 for each donor. Cells were banked down at P3 and all *in vitro* experiments were carried out between P5-7.

#### Immunophenotype Characterization

ECFCs were passaged and then identified using flow cytometry by staining 100,000 cells with 2.5 µg/ml CD31-FITC (Biolegend, 303104), 0.5 µg/ml CD34-APC (Biolegend, 342510), 1.25 µg/ml VEGFR2-PE (BD, 560494), and 0.06 µg/ml CD45-PE-Cy7 (Biolegend, 368532) diluted in flow cytometry buffer (FCB) (2% FBS in PBS) for 20 mins at 4^o^C in the dark. To gate out dead cells 0.15 µM DRAQ7 (Thermofisher, D15106) in FCB was added 90 sec prior to running a minimum of 20,000 cell events on a Canto II flow cytometer (BD). Data analysis was carried out on Flowjo X (v10.0.7r2).

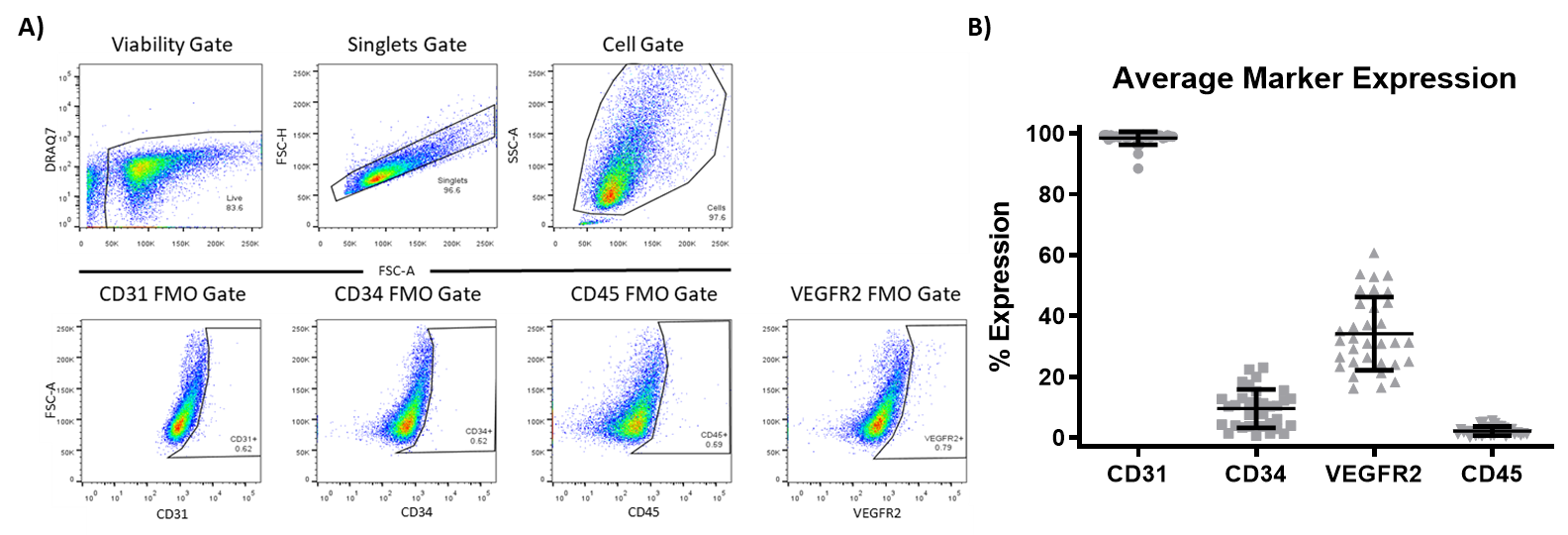

Supplemental Figure S1. Isolated cells were identified as ECFCs.

*A) Gating strategy. B) ECFC phenotypic characterization with the percentage of positive cells expressing each marker indicated. N = 33. FMO = Fluorescence Minus One. Mean ± SD.*

Supplementary Table S1. Cell number needed for an ECFC therapeutic.

| Experiment | Number of ECFCs needed | Spare ECFCs | Total |
| --- | --- | --- | --- |
| Flow Cytometry Cell Characterisation | 0.7x10^6^ | 0.1x10^6^ | 0.8x10^6^ |
| Karyotype analysis | 1x10^6^ | 1x10^6^ | 2x10^6^ |
| Matrigel | 0.066x10^6^ | 0.022x10^6^ | 0.088x10^6^ |
| Sterility, Endotoxin, Mycoplasma (8% of final product) | 35.2x10^6^ – 87x10^6^ | 35.2x10^6^ – 87x10^6^ | 70.4x10^6^– 174x10^6^ |
| Transplantation (0.1x10^6^ – 2.5x10^6^ $\sim$20 g mouse) | 0.44x10^9^ – 10.88x10^9^ in 87 kg human | 0.04x10^9^ – 1.09x10^9^ | 0.48x10^9^-11.96x10^9^ |
| **Total number of cells needed** |  |  | **0.55x10^9^ – 12.13x10^9^** |

Supplementary Table S2. Predictors of the ability to isolate ECFCs from the PB of patients.

| Parameter | No ECFCs Isolated | ECFCs Isolated | P-value |
| --- | --- | --- | --- |
| **ECFCs per 10^8^ PBMCs** | **2.34 ± 3.11** | **21.43 ± 42.28** | **<0.0001** |
| **Creatinine (mmol/L)** | **69.76 ± 21.54** | **82.45 ± 17.44** | **0.007** |
| **High Density Lipoproteins (mmol/L)** | **1.51 ± 0.62** | **1.15 ± 0.36** | **0.03** |
| **Weight (kg)** | **76.86 ± 15.57** | **87.54 ± 15.59** | **0.03** |
| **Mean Corpuscular Haemoglobin (pg)** | **31.08 ± 1.69** | **29.72 ± 2.36** | **0.03** |
| **Eosinophils (10^9^/L)** | **0.18 ± 0.19** | **0.27 ± 0.15** | **0.03** |
| **Mean Corpuscular Volume (fl)** | **92.25 ± 4.35** | **88.53 ± 6.31** | **0.04** |
| **Systolic Pressure (mm Hg)** | **123.80 ± 12.32** | **134.40 ± 17.02** | **0.046** |
| Body Mass Index (kg/m^2^) | 26.58 ± 5.22 | 29.75 ± 6.03 | 0.07 |
| HbA1c (mmol/mol) | 47.29 ± 16.05 | 53.21 ± 14.76 | 0.07 |
| Diastolic Pressure (mm Hg) | 70.21 ± 7.77 | 74.93 ± 7.93 | 0.08 |
| Potassium (mmol/L) | 4.27 ± 0.46 | 4.45 ± 0.36 | 0.13 |
| Isolated Blood Volume (ml) | 84.08 ± 12.05 | 77.18 ± 16.04 | 0.13 |
| Fibrinogen (g/L) | 4.13 ± 0.85 | 3.78 ± 1.29 | 0.13 |
| Total Protein (g/L) | 73.94 ± 4.37 | 71.3 ± 6.57 | 0.14 |
| Cholesterol/High Density Lipoprotein Ratio | 3.22 ± 1.41 | 3.61 ± 1.81 | 0.17 |
| Triglycerides (mmol/L) | 1.57 ± 0.85 | 1.96 ± 1.06 | 0.18 |
| Monocytes (10^9^/L) | 0.55 ± 0.34 | 0.62 ± 0.20 | 0.19 |
| Gamma-Glutamyltransferase (U/L) | 63.94 ± 73.95 | 39.27 ± 35.56 | 0.22 |
| Alanine Transaminase (U/L) | 29.29 ± 14.81 | 25.79 ± 15.41 | 0.23 |
| Albumin (g/L) | 46.06 ± 4.09 | 44.67 ± 3.93 | 0.27 |
| C-Peptide (pmol/L) | 1810 ± 1057 | 1703 ± 1527 | 0.37 |
| Ankle Brachial Index | 0.79 ± 0.63 | 0.60 ± 0.33 | 0.39 |
| Cholesterol (mmol/L) | 4.29 ± 1.08 | 4.05 ± 1.10 | 0.45 |
| Chloride (mmol/L) | 101.4 ± 2.06 | 101.5 ± 4.62 | 0.45 |
| Height (cm) | 170.10 ± 9.60 | 172.10 ± 8.62 | 0.46 |
| Red Blood cells (10^12^/L) | 4.78 ± 1.69 | 4.63 ± 0.62 | 0.49 |
| Haematocrit (L/L) | 0.40 ± 0.05 | 0.41 ± 0.05 | 0.50 |
| White Cell Count (10^9^/L) | 7.71 ± 2.52 | 7.71 ± 1.982 | 0.53 |
| Alkaline Phosphatase (U/L) | 77 ± 21.17 | 85.73 ± 31.65 | 0.56 |
| Mean Corpuscular Haemoglobin Concentration (g/dl) | 33.73 ± 0.84 | 33.56 ± 1.00 | 0.56 |
| Random Glucose (mmol/L) | 6.81 ± 1.93 | 8.82 ± 4.57 | 0.57 |
| Basophils (10^9^/L) | 0.05 ± 0.03 | 0.05 ± 0.04 | 0.57 |
| International Normalised Ratio | 1.01 ± 0.09 | 1.09 ± 0.48 | 0.60 |
| Sodium (mmol/L) | 140.4 ± 2.18 | 139.2 ± 4.37 | 0.61 |
| Estimate Glomerular Filtration Rate (ml/min) | 99.75 ± 20.45 | 98.03 ± 21.37 | 0.62 |
| Neutrophils (10^9^/L) | 4.93 ± 2.07 | 4.81 ± 1.67 | 0.65 |
| Age (yea rs) | 64.41 ± 9.11 | 65.55 ± 8.48 | 0.66 |
| Total Bilirubin (μmol/L) | 7.71 ± 3.87 | 7.88 ± 3.44 | 0.66 |
| Adjusted Calcium (mmol/L) | 2.37 ± 0.10 | 2.39 ± 0.11 | 0.67 |
| Diabetes Duration (years) | 8.86 ± 4.10 | 10.08 ± 6.86 | 0.67 |
| Lymphocytes (10^9^/L) | 1.91 ± 0.63 | 1.98 ± 0.69 | 0.73 |
| Total Calcium (mmol/L) | 2.39 ± 0.14 | 2.39 ± 0.13 | 0.77 |
| Phosphorous (mmol/L) | 1.06 ± 0.21 | 1.07 ± 0.17 | 0.78 |
| Lactate Dehydrogenase (U/L) | 199 ± 32.64 | 195.6 ± 42.75 | 0.78 |
| Activated Partial Thromboplastin Time (s) | 26.31 ± 5.81 | 25.83 ± 3.38 | 0.81 |
| Urea (mmol/L) | 5.78 ± 1.94 | 5.9 ± 1.79 | 0.83 |
| Haemoglobin (g/dL) | 13.43 ± 1.87 | 13.39 ± 2.03 | 0.86 |
| Prothrombin time (s) | 10.88 ± 0.86 | 11.60 ± 4.60 | 0.89 |
| Low Density Lipoprotein (mmol/L) | 2.08 ± 1.07 | 2.08 ± 1.01 | 0.98 |
| Platelet (10^9^/L) | 246.2 ± 83.06 | 270.2 ± 91.34 | >0.99 |
| # of isolated PBMCs | 183x10^6^ ± 0.72x10^6^ | 195x10^6^ ± 0.96x10^6^ | 0.99 |

*Patients were divided by the ability to successfully isolate ECFCs and then the clinical and biochemical characteristics were compared between groups. Data presented as mean ± SD and ranked by p-value.*
