## Supplementary Table 1 for "An Assessment of the Functional State of Endothelial Colony Forming Cells from Patients with Diabetes Mellitus and Chronic Limb Threatening Ischemia"

| Parameter | AMC | Diabetic | CLTI | Diabetic-CLTI | p-value |
| --- | --- | --- | --- | --- | --- |
| Patient number | 8 | 9 | 6 | 10 | - |
| **Patient Characteristics** | | | | | |
| Age (years) | 60.38 (9.9) | 65.33 (9.77) | 68.83 (6.56) | 68 (6.94) | 0.20 |
| BMI (kg/m^2^) | 31.05 (6.75) | 30.53 (4.81) | 26.68 (5.46) | 29.55 (6.98) | 0.63 |
| Gender (M/F) | 4/4 | 7/2 | 4/2 | 9/1 | n.s. |
| Smoking (Yes/Past/ Never/unknown) | 3/2/3/0 | 0/4/4/1 | 2/2/2/0 | 1/7/2/0 | - |
| **Clinical Characteristics** | | | | | |
| **ABI (lowest value)** | **N/A** | **N/A** | **0.44 (0.23)** | **0.96 (0.53)†** | **0.04** |
| Diabetic Duration (years) | N/A | 13.17 (7.47) | N/A | 7.77 (5.77) | 0.15 |
| Hypertensive (Yes/no/unknown) | 1/6/1 | 1/9/0 | 2/3/1 | 4/5/1 | 0.15 |
| Diastolic Pressure (mm Hg) | 75.7 (9.7) | 75.5 (7.6) | 76.8 (5.9) | 74.0 (9.0) | 0.92 |
| Systolic Pressure (mm Hg) | 144.3 (13.7) | 136.5 (19.3) | 128.5 (23.9) | 131.1 (15.0) | 0.44 |
| **Blood Biochemistry** | | | | | |
| aPTT (s) | 24.99 (2.25) | 26.31 (4.66) | 24.32 (2.02) | 27.02 (3.49) | 0.28 |
| ALT (U/L) | 28.5 (14.8) | 29.2 (18.2) | 14.5 (3.8) | 27.3 (16.3) | 0.27 |
| C-Peptide (pmol/L) | 1670.4 (1676.0) | 2095.7 (1531.99) | 2428.5 (2015.65) | 941.9 (780.64) | 0.13 |
| **Cholesterol (mmol/L)** | **5 (1.17)** | **3.54 (0.68)*** | **3.58 (0.98)** | **3.97 (1)** | **0.02** |
| Creatinine (mmol/L) | 73.5 (1.4) | 91.6 (11.6) | 83.5 (17.3) | 80.8 (21.9) | 0.16 |
| eGFR (ml/min) | 107.6 (22.3) | 90.6 (19.8) | 96 (20.5) | 98.1 (22.7) | 0.53 |
| **HbA1c (mmol/mol)** | **38.9 (2.2)** | **57.0 (12.4)*** | **43.0 (5.0)** | **67.4 (5.0)*†** | **<0.0001** |
| HDLs (mmol/L) | 1.14 (0.4) | 1.14 (0.32) | 1.15 (0.31) | 1.17 (0.44) | 0.998 |
| LDLs (mmol/L) | 2.95 (1.23) | 1.81 (0.43) | 1.7 (0.81) | 1.81 (0.97) | 0.04 |
| Platelets (10^9^/L) | 222.4 (22.1) | 250.1 (53.3) | 317.2 (156.2) | 296.4 (92.1) | 0.12 |
| PT (s) | 10.78 (0.71) | 14.01 (9.15) | 10.67 (0.33) | 10.88 (0.57) | 0.77 |
| **Random Glucose (mmol/L)** | **5.5 (1.3)** | **9.2 (2.9)** | **5.9 (1.3)** | **11.7 (5.9)*** | **0.04** |
| Triglycerides (mmol/L) | 2.79 (1.06) | 1.64 (0.94) | 1.58 (0.4) | 1.8 (1.2) | 0.06 |
| Urea (mmol/L) | 5.81 (1.44) | 6.52 (1.42) | 4.37 (1.39) | 6.33 (2.19) | 0.1 |
| **Medications** | | | | | |
| Ace Inhibitors (yes/no) | 0/8 | 3/6 | 3/3 | 1/9 | n.s. |
| β-Blockers (yes/no) | 0/8 | 1/8 | 3/3 | 4/6 | n.s. |
| **Biguanides (yes/no)** | **0/8** | **9/0*** | **1/5** | **6/4*** | **0.003 0.0007** |
| Factor Xa Inhibitors (yes/no) | 1/7 | 1/8 | 1/5 | 3/7 | n.s. |
| **NSAIDs (yes/no)** | **0/8** | **4/5** | **4/2*** | **7/3*** | **0.02 0.004** |
| PPI (yes/no) | 0/8 | 2/7 | 1/5 | 2/8 | n.s. |
| SGLT2 Inhibitors (yes/no) | 0/8 | 1/8 | 0/6 | 2/8 | n.s. |
| **Statins (yes/no)** | **2/6** | **2/7** | **2/4** | **8/2^$^** | **0.02** |
