## Supplementary Table 2 for "An Assessment of the Functional State of Endothelial Colony Forming Cells from Patients with Diabetes Mellitus and Chronic Limb Threatening Ischemia"

|  | Step 1: Individual Variables | | | Step 2: Multivariate | | | Step 3: Refined Multivariate | | |
| --- | --- | --- | --- | --- | --- | --- | --- | --- | --- |
| **Risk Factor** | **Odds ratio** | **95% CI** | **R^2^ (adj) (%)** | **Odds ratio** | **95% CI** | **R^2^ (adj) (%)** | **Odds ratio** | **95% CI** | **R^2^ (adj) (%)** |
| Age | 1.02 | (0.95, 1.10) | 0 |  |  | 65.17 |  |  | 68.43 |
| BMI | 1.13 | (0.99, 1.28) | 5.28 |  |  |  |  |  |  |
| Chol/HDL Ratio | 1.34 | (0.77, 2.33) | 0.24 |  |  |  |  |  |  |
| Cholesterol | 0.72 | (0.40, 1.29) | 0.45 |  |  |  |  |  |  |
| C-Peptide | 1.00 | (1.00, 1.00) | 0 |  |  |  |  |  |  |
| Creatinine | **1.04** | **(1.00, 1.08)** | **8.02** | 1.03 | (0.94, 1.13) |  |  |  |  |
| ECFCs/10^8^ PBMCs | **1.91** | **(1.11, 3.28)** | **38.51** | **4.34** | **(1.04, 18.14)** |  | **4.04** | **(1.22, 13.38)** |  |
| HbA1c | 1.03 | (0.98, 1.08) | 1.02 |  |  |  |  |  |  |
| HDL | **0.18** | **(0.04, 0.73)** | **9.84** | 0.05 | (0.0004, 5.83) |  |  |  |  |
| LDL | 0.89 | (0.49, 1.62) | 0 |  |  |  |  |  |  |
| Random Glucose | 1.16 | (0.94, 1.43) | 2.67 |  |  |  |  |  |  |
| Systolic Pressure | **1.06** | **(1.00, 1.12)** | **8.59** | 1.20 | (1.00, 1.44) |  | **1.18** | **(1.03, 1.35)** |  |
| Triglycerides | 1.76 | (0.80, 3.85) | 2.42 |  |  |  |  |  |  |
